# Insulin signaling mediates neurodegeneration in glioma

**DOI:** 10.1101/2020.01.03.894469

**Authors:** Patricia Jarabo, Carmen de Pablo, Héctor Herranz, Francisco Antonio Martín, Sergio Casas-Tintó

## Abstract

Cell to cell communication facilitates tissue development and physiology. Under pathological conditions, brain tumors disrupt glia-neuron communication signals that in consequence, promote tumor expansion at the expense of surrounding healthy tissue. The glioblastoma is the most aggressive and frequent brain tumor. This type of glioma expands and infiltrates into the brain, causing neuronal degeneration and neurological decay, among other symptoms. Here we describe how the glioblastoma produce ImpL2, an antagonist of the insulin pathway, which is regulated by the microRNA *miR-8*. ImpL2 targets neighboring neurons and causes mitochondrial disruption as well as synapse loss, both early symptoms of neurodegeneration. Furthermore, glioblastoma progression requires insulin pathway attenuation in neurons. Restoration of neuronal insulin activity is sufficient to rescue the synapse loss and to delay the premature death caused by glioma. Therefore, signals from GB to neuron emerge as a potential field of study to prevent neurodegeneration and to develop anti-tumoral strategies.

**Significance Statement:** Glioblastoma (GB) is the most aggressive type of brain tumour and currently there is no cure or effective treatment. Patients course with neurological decay and previous data in animal models indicate that GB cause a neurodegenerative process. We describe here a molecule named ImpL2 that is produced by GB cells and impact on neighbouring neurons. ImpL2 is an antagonist of the insulin pathway and signaling reduction in neurons causes mitochondrial defects and synapse loss. These mechanisms underlying GB-induced neurodegeneration plays a central role in the premature death caused by this tumour. Restoration of insulin signaling in neurons prevents tumour progression and rescues the lethality caused by GB in *Drosophila* models.

## Introduction

Cancer is one of the leading causes of mortality worldwide and is expected to be responsible for 15 million deaths in 2020 (65% in less developed countries) according to the World Health Organization. Notwithstanding recent advances in health treatments and extended lifespan of patients, some tumors still remain incurable. Among them, glioblastoma (GB) stands out because it is the most frequent and aggressive primary brain tumor. It is originated from glial cells and causes death within the first year after diagnosis (1), despite standard treatments such as resection, radiotherapy, and chemotherapy. This is accompanied by broad neurological dysfunctions (2). Brain tumors cause cognitive decline and neuronal dysfunction (reviewed in 3, 4). These cognitive defects are consistent with typical neurodegenerative-associated symptoms such as synapse loss and mitochondrial alterations (5, 6).

*Drosophila melanogaster*, the fruit fly, has emerged as a reliable animal system to mimic human diseases such as cancer (7). However, the aim is not to heal a sick insect but to model cellular and molecular mechanisms of human diseases, in order to identify targets for eventual diagnosis and treatments of patients. The power of *Drosophila* genetics allows genetic and pharmacological screens that may be translated to medicine, particularly for neurodegenerative disorders (8-13). In fact, a *Drosophila* GB model that recapitulates most of the human disease features has been developed and validated (10, 14-16). This model is based on two of the most frequent mutations in patients, a constitutively active form of the epidermal growth factor receptor (dEGFRλ) and the phosphatidylinositol-3 kinase (PI3K) catalytic subunit p110α (PI3K92E) driven by the glial specific *repo-Gal4* (16). This animal model has brought novel understandings into GB molecular mechanisms (10, 14, 17, 18).

MicroRNAs (miRNAs) are short non-coding RNAs that control gene activity mainly through post-transcriptional mechanisms. Recently, they have been related to almost all biological processes and diseases, particularly cancer (19, 20). Related to glioma, the *miR-200* family (which includes *miR-200, miR-141* and *miR-429*) plays central roles in GB development, metastasis, therapeutic response, and prognosis (reviewed by (21)). Low levels of *miR-200* are indicative of poor prognosis in GB (22). In colorectal cancer and GB, low expression levels of mi-RNAs correlates with up-regulation of *insulin-like growth binding protein 7* (*IGFBP7*) (23). Likewise, in GB there is a Transforming Growth Factor Beta 2 (TGFB2)-dependent increase in IGFBP7 protein levels (24). However, the mechanisms involved in IGFBP7 influence on GB progression and its regulation by *miR-200* remains unsolved. In *Drosophila, miR*-*200* and *IGFBP7* are represented by *miR-8* and *Imaginal morphogenesis protein-late 2* (*ImpL2*), respectively (25). In juvenile stages, *miR-8* has been related to glial cell growth and positively regulates positively synaptic growth at the neuromuscular junction (26, 27).

In contrast, *Drosophila ImpL2* is related to cachexia, a systemic effect characterized by anorexia and metabolic alterations induced by other malignant tumors (28). Secreted ImpL2 from epithelial tumor cells induces systemic organ wasting and insulin resistance by antagonizing insulin signaling (29, 30). Interestingly, PI3K and *Drosophila Ras homolog enriched in brain* (dRheb), two members of the insulin pathway, induce the formation of synapsis between neurons (a process known as synaptogenesis) in the *Drosophila* larval brain (31). Actually, it has been shown that AKT, also involved in insulin signaling, acts as a pro-synaptogenic element (32). These data strongly support a role for insulin signaling in the regulation of neuronal synaptic activity in *Drosophila*. In mammals, a similar effect of insulin signaling on synaptic plasticity has been described (33). Notably, synapse loss is an early step in neurodegeneration (34, 35). We have recently re-evaluated GB as a neurodegenerative disease, showing that GB reduces the number of synapsis through wingless/frizzled 1 (wg/fz1) signaling (Portela et al, PLOS Biol 2019), equivalent to mammalian WNT pathway (36). However, whether tumoral glial cells are able to modify insulin signaling directly in neurons, and consequently alter the number of synapses, is yet unknown.

In this report, we show that secreted ImpL2 from glial-derived tumoral cells antagonizes insulin signaling in neighboring neurons, inducing a reduction in synapse number and consequently promoting neurodegeneration. *ImpL2* expression in GB cells is regulated by *miR-8*, thus linking functionally micro-RNA pathway with Insulin signaling in a GB model. We describe the function of ImpL2 as a mediator in GB-neuron communication, responsible for the reduction in synapse number and neurological defects. Indeed, we propose the insulin pathway as a core signal in GB progression and neurological decay. Finally, we propose a novel neuroprotective strategy against GB that extend lifespan and improve life quality.

## Results

### ImpL2 mediates GB progression and neurodegeneration

To study the mechanisms of communication among malignant glial cells and neurons, we used a previously well-characterized *Drosophila* GB model that stimulates the oncogenic transformation of glial cells and lethal glial neoplasia in post-embryonic larval (14, 16) or adult brains (10), leading to lethal glial neoplasia. We previously reported a reduction in the number of synapses in the neuromuscular junction (NMJ) of adult flies caused by GB progression (Portela et al, 2019). This neurodegenerative process is observed even though those genetic modifications were caused in glial cells. This phenomenon suggests that signals originated in the glial tumor impact on neighboring neurons. Synaptogenesis is tightly regulated by PI3K, a main player in insulin signaling pathway (32). Moreover, the expression of secreted molecules that decrease insulin pathway activity, such as ImpL2 (IGFBP7 in humans), correlates with GB progression (23, 24).

To determine *ImpL2* mRNA expression levels in GB we performed qPCR experiments. *ImpL2* mRNA shows an increase in GB samples as compared to control brains (fig 1A). To discriminate *ImpL2* expression in neuronal or glial (GB) cells, we used a MIMIC GFP reporter that reproduces faithfully *ImpL2* expression (37). Consistently, GB cells show higher GFP levels than control glial cells. Likewise, upon *ImpL2* RNAi expression we detect a decrease in GFP levels, similar to the ones observed in control brains (fig 1B-D). In a previous work we established that tumoral progression depends on the formation of a network of protrusions (i.e. an expansion of the membrane surface) named tumor microtubules (TMs), similarly to human GB (17, 38). Besides, in mammals and flies the TM network requires the *GAP43* and *igloo* gene functions, respectively (17, 38). We also showed that GB progression requires c-Jun N-terminal Kinase (JNK) pathway activity (17). The *Drosophila* JNK homolog *basket* (*bsk*) plays a central role in JNK signaling in normal and tumoral conditions (39). We overexpressed *igloo RNAi* and a dominant negative form of *bsk* to block TM formation and JNK activity, respectively. As expected, *ImpL2* expression levels are reverted to similar levels to the ones observed in controls brains in both cases (fig 1E-F). Indeed, the analysis of co-localization rate between ImpL2 and glial cells shows a significant increase in ImpL2 levels specifically in GB cells (fig 1G).

**Fig 1.**
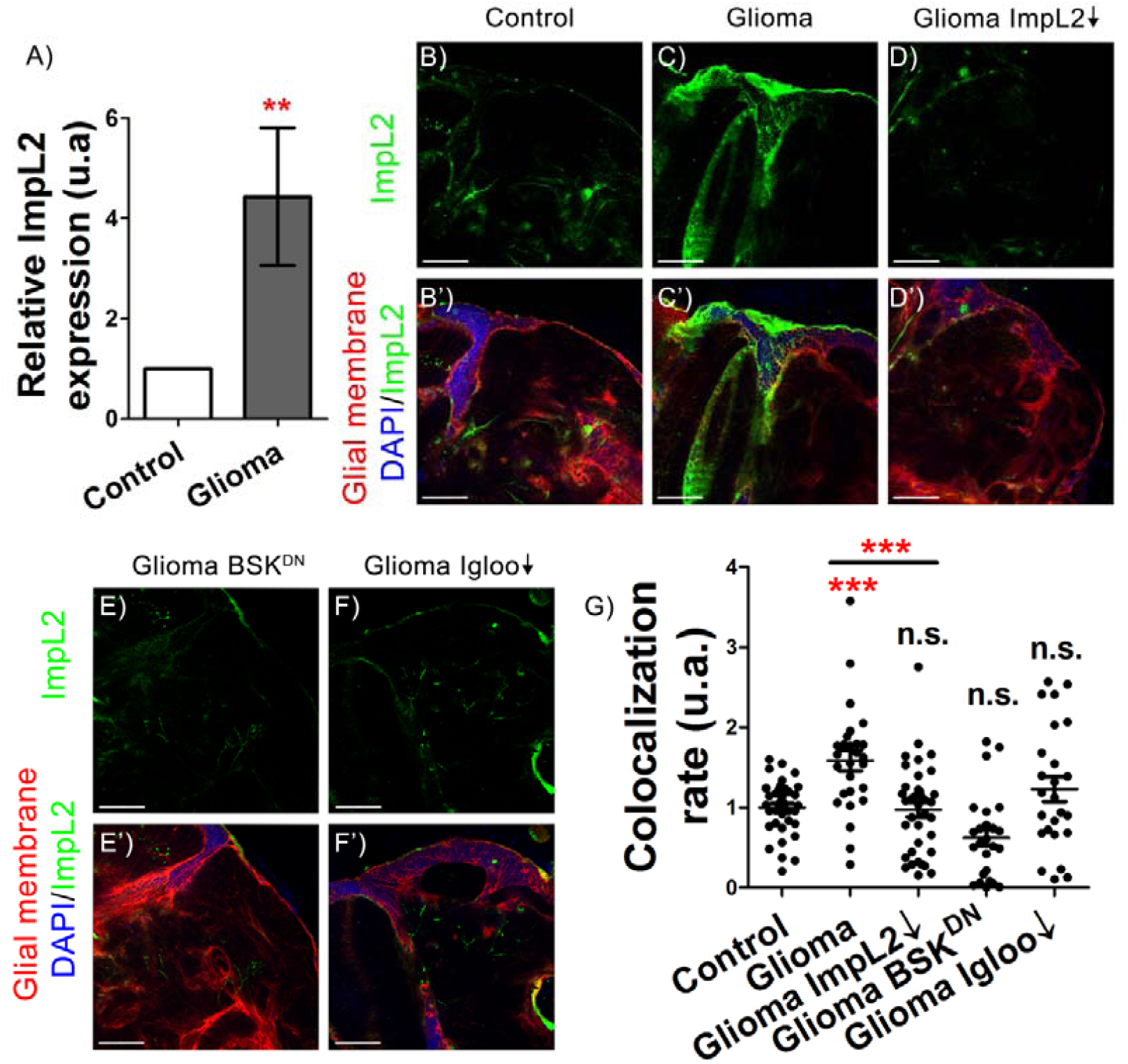
ImpL2 is upregulated in gliomas. A) RT-qPCR shows an upregulation of *ImpL2* in gliomas of repo> *UAS-dEGFR*^λ^, *UAS-dp110*^*CAAX*^ flies (t-test) in at least *N* = 30 per genotype. Confocal microscopy images of 7 day old adult brains from B) *repo>UAS-LacZ* (Control), C) *repo>UAS-dEGFR*^*λ*^, *UAS-dp110CAAX* (Glioma), D) *repo>UAS-dEGFR*^λ^, *UAS-dp110CAAX*; *UAS-Impl2 RNAi* (*Glioma>ImpL2↓*), E) *repo>UAS-dEGFR*^λ^, *UAS-dp110CAAX*; *UAS-BSK*^*DN*^ (*Glioma>BSK*^*DN*^) and F) *repo>UAS-dEGFR*^λ^, *UAS-dp110CAAX*; *UAS-igloo RNAi* (*Glioma>igloo↓*) animals, in all cases together with an *ImpL2-MIMIC GFP* transgene G) Quantification and statistical analysis of correlation rate between *Impl2-MIMIC GFP* and glial membrane in at least *N* = 10 per genotype (ANOVA, post-hoc Bonferroni) (scale bar, 50 μm) (\*\**p*-value>0,005, \*\*\**p*-value>0,0001).

To detect signs of neurodegeneration, we quantify the number of synapses in the neuromuscular junction (NMJ) of the adult flies with GB. NMJ is a stereotyped structure that allows to count the number of synapses (i.e.synapsessynapses) in the synaptic buttons of the motoneurons by using anti-bruchpilot, an specific antibody that recognizes synapses unambiguously (see material and methods for details). To determine the contribution of *ImpL2* to synapse loss, we knockdown *ImpL2* in GB cells and counted the number of synapsessynapses in adult NMJs. The results show that *ImpL2* reduction in GB cells counteracted the reduction in the number of synapsessynapses of GB brains (fig 2A-C, E). Consistently, *ImpL2* over-expression in wild type (wt) glial cells decreases the number of synapse in NMJs (fig 2D-E). Two additional and typical features of GB are the increase in the number of glial cells and the expansion of the TM network (17) (Fig 2F-G, I-J). As expected, the down-regulation of ImpL2 RNA levels in GB causes a striking reduction in the number of glial cells (fig 2H-I) and in the total tumor volume (2J).

**Fig 2.**
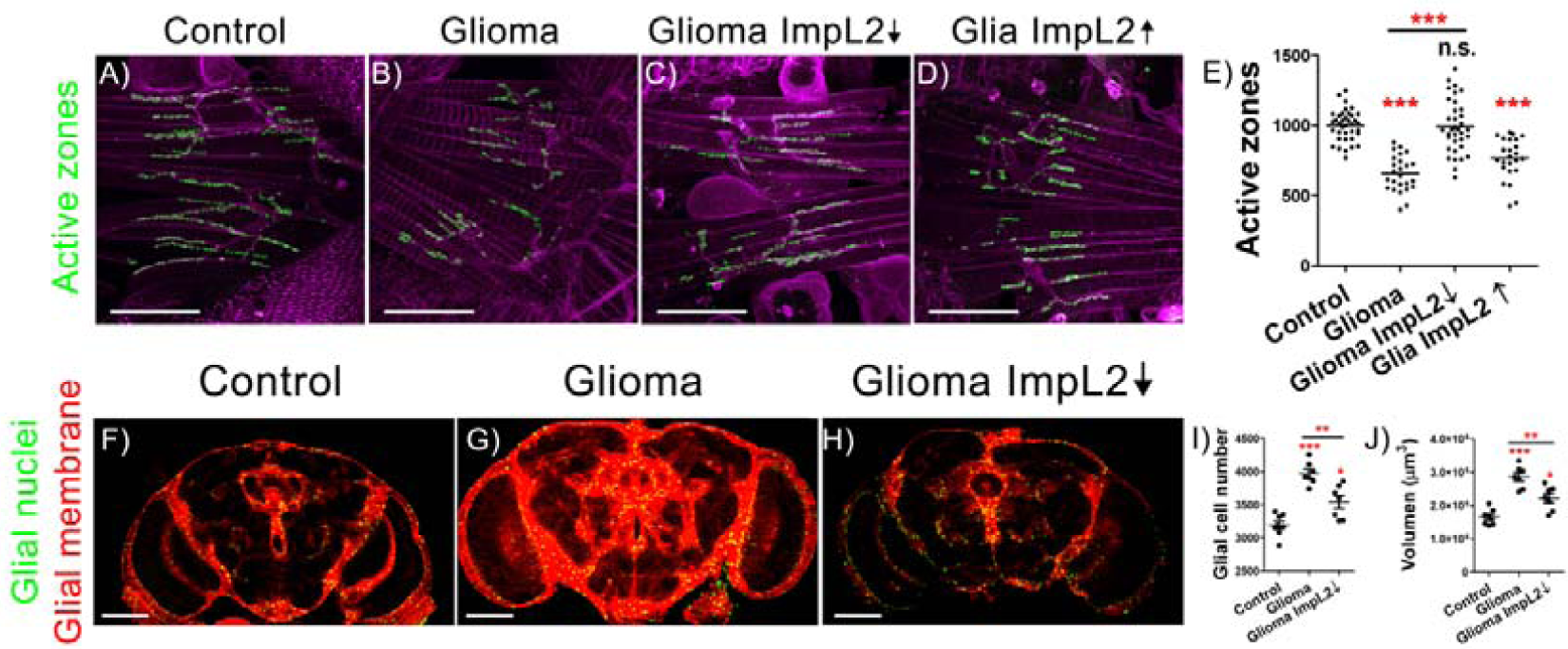
ImpL2 downregulation in glioma cells causes neurodegeneration and reduces tumor progression. Confocal Images of adult neuromuscular junction (NMJ) at 7th day at 29ºC from A) *repo>LacZ*, B) *repo>UAS-dEGFR*^λ^, *UAS-dp110CAAX*, C) *repo>UAS-dEGFR*^λ^, *UAS-dp110CAAX*; *UAS-Impl2 RNAi* and D) *repo>UAS-dEGFR*^λ^, *UAS-dp110CAAX*; UAS-*ImpL2* animals. Active zones are marked in green (anti-NC82). E) Quantification and statistical analysis of active zones in at least *N* = 13 per genotype (ANOVA, post-hoc Bonferroni) (scale bar, 50 μm). Confocal microscopy images of adult brains from F) *repo>UAS-LacZ*, G) *repo>UAS-dEGFR*^λ^, *UAS-dp110CAAX* and H) *repo>UAS-dEGFR*^λ^, *UAS-dp110CAAX*; *UAS-Impl2 RNAi* flies after 7 days at 29ºC. Glial membrane is in red and glial nuclei in green. Quantification of I) glial cell number and J) glial membrane volume for at least N = 7 per genotype (ANOVA, post-hoc Bonferroni) (scale bar, 100 μm). (\**p*-value>0,05\*\**p*-value>0,005, \*\*\**p*-value>0,0001).

Thus, we conclude that GB cells upregulate and secrete ImpL2, a necessary step to induce tumoral expansion and a reduction of the synapse number in surrounding neurons.

### microRNAs regulate *ImpL2* expression in GB

It has been described that low levels of micro-RNAs correlate with high levels of ImpL2 homolog in human GB, suggesting that ImpL2 regulation might be mediated by microRNAs (23). Accordingly, there is a down-regulation of the microRNA *miR-200* family in GB samples (reviewed in (21)) Given that *miR-8*, the Drosophila homologue of *miR-200* family, negatively regulates *ImpL2* mRNA stability in the fat body (40), we hypothesize that *miR-8* may play a role in GB progression by regulating ImpL2 levels. We used a *miR-8* sensor to monitor *miR-8* activity. It includes *miR-8* binding sites in the 3’UTR of the *GFP* gene (41). Thus, high levels of GFP indicate low levels of *miR-8* activity, and vice versa (see Materials and Methods). GFP signal is increased in GB cells (fig 3A-C), indicating that *miR-8* levels are reduced. To determine if the increase of *ImpL2* in GB depend on *miR-8*, we analyzed *ImpL2* sensor upon *miR-8* overexpression in that context. We observed a significant reduction of *ImpL2* expression in GB cells *in vivo*. Consistently, *miR-8* gain-of-function in GB partially rescues the loss of synapses, recapitulating the effect of *ImpL2* loss-of-function in GB conditions (fig 3D-G). In addition, the reduction of synapse number in NMJ caused by the GB can be attenuated by *miR-8* overexpression in GB cells (fig 3H-J, L). This is consistent with the effect of *ImpL2* down-regulation in GB that also rescued the synapse number (fig 2C). Furthermore, *miR-8* overexpression in wt glial cells (with low *ImpL2* levels) does not alter the number of synapses (fig 3K-L). Intriguingly, GB cell number expansion is not prevented by *miR-8* overexpression (fig 3M-O, Q), and consistently *miR-8* overexpression does not change glial cell number in wild type conditions (fig 3P-Q). However, we do observe a reduction of GB membrane volume upon *miR-8* upregulation, something that does not occur in normal glial cells (fig 3R). Altogether these results show an inverse correlation between miR-8 and ImpL2 expression in GB cells and suggest that ImpL2 levels are regulated by *miR-8 in vivo*.

**Fig 3.**
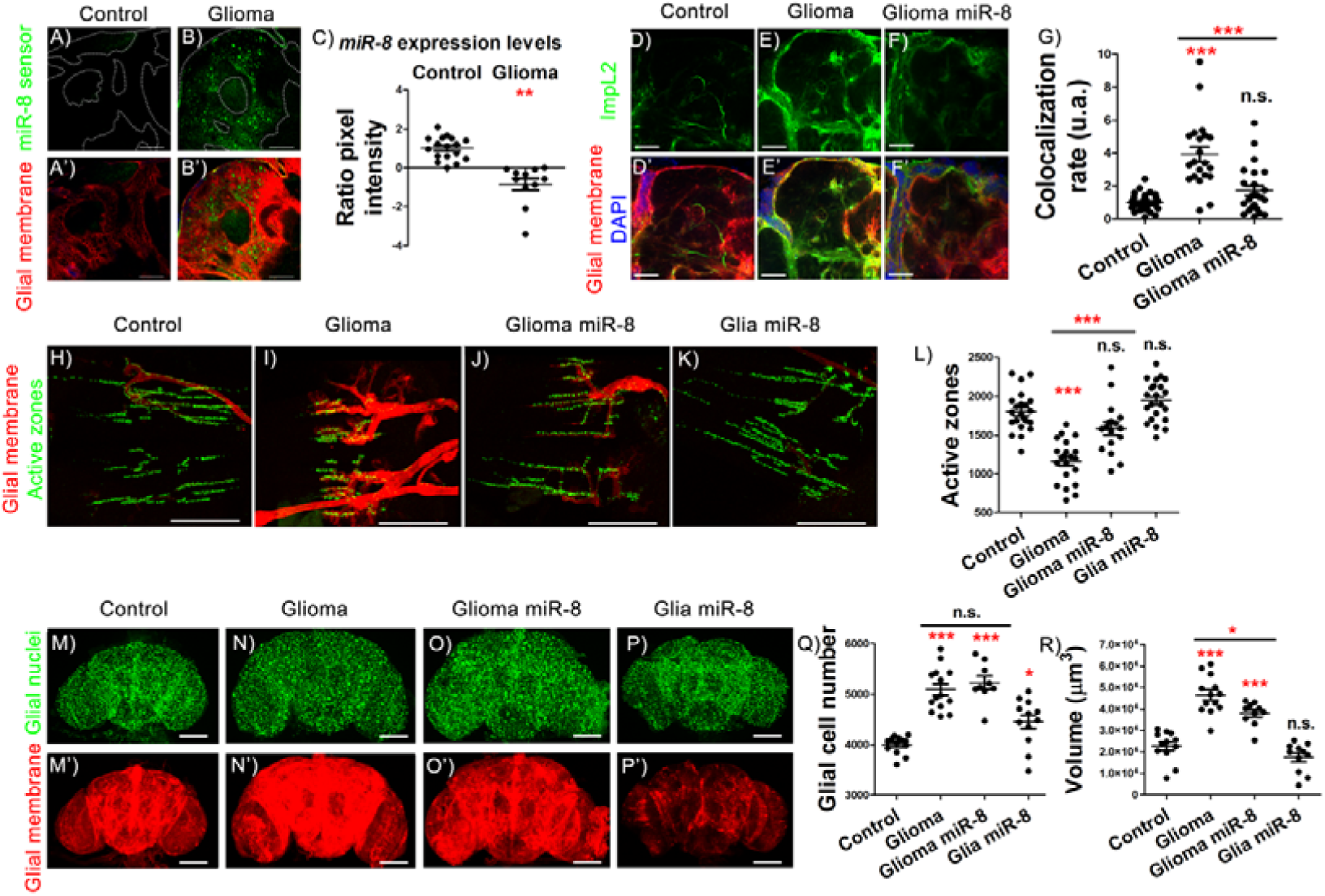
Increasing *miR-8* levels in gliomas impairs ImpL2 upregulation and prevents synapse loss. A-B) Confocal microscopy images of adult brain showing the expression pattern of *miR-8* in *repo>UAS-LacZ* control animals compare to *repo>UAS-dEGFR*^λ^, *UAS-dp110CAAX* flies using *miR-8* sensor (see Materials and Methods). A’-B’) Glial membrane is marked in red. C) Quantification and statistical analysis of pixel intensity in at least *N* = 3 per genotype (ANOVA, post-hoc Bonferroni) (scale bar, 50 mm). Confocal microscopy images of adult brains from D) *repo>UAS-LacZ*, E) *repo>UAS-dEGFR*^λ^, *UAS-dp110CAAX*, F) *repo>UAS-dEGFR*^λ^, *UAS-dp110CAAX*; *UAS-miR-8* using an *ImpL2-MIMIC* line after 7 days at 29ºC. G) Quantification and statistical analysis of correlation rate in at least *N* = 9 per genotype (ANOVA, post-hoc Bonferroni) (scale bar, 50 μm). Adult neuromuscular junction (NMJ) from H) *repo>UAS-LacZ*, I) *repo>UAS-dEGFR*^λ^, *UAS-dp110CAAX*, J) *repo>UAS-dEGFR*^λ^, *UAS-dp110CAAX*; *UAS-miR-8* and K) *repo>miR-8 after 7 days at 29ºC*. Active zones are marked in green. L) Quantification and statistical analysis of active zones number in at least *N* = 10 per genotype (ANOVA, post-hoc Bonferroni) (scale bar, 50 mm). Confocal microscopy images of adult brain from 7-day-old flies of M) *repo>UAS-LacZ*, N) *repo>UAS-dEGFR*^λ^, *UAS-dp110CAAX*, O) *repo>UAS-dEGFR*^λ^, *UAS-dp110CAAX, UAS-miR-8* and P) *repo>UAS-LacZ, UAS-miR-8* genotypes. M’-P’) Glial membrane is shown in red and glial nuclei in green. Quantification of Q) glial cells and R) glial membrane volume for at least N = 9 per genotype (ANOVA, post-hoc Bonferroni) (scale bar, 100 mm). (\**p*-value>0,05,\*\**p*-value>0,005, \*\*\**p*-value>0,0001)

### GB secreted ImpL2 reduces neuronal Insulin signaling

To evaluate the impact of Insulin signaling reduction in neurons, we measure *dRheb* mRNA by qPCR. d*Rheb* is the molecular link between the insulin signaling pathway and TOR kinase, and it reflects the Insulin pathway activity (reviewed in (42)): d*Rheb mRNA* levels drop down when Insulin signaling is low (43). We analyzed d*Rheb* expression levels in control and GB brains by qPCR. The GB itself is induced by overexpressing a constitutively active form of PI3K, thus the insulin pathway is activated in all glial cells. However, mRNA levels of *dRheb* are reduced in GB brains when compared with control brains, suggesting that this increase reflects mostly neuronal expression (fig 4E).

**Fig 4.**
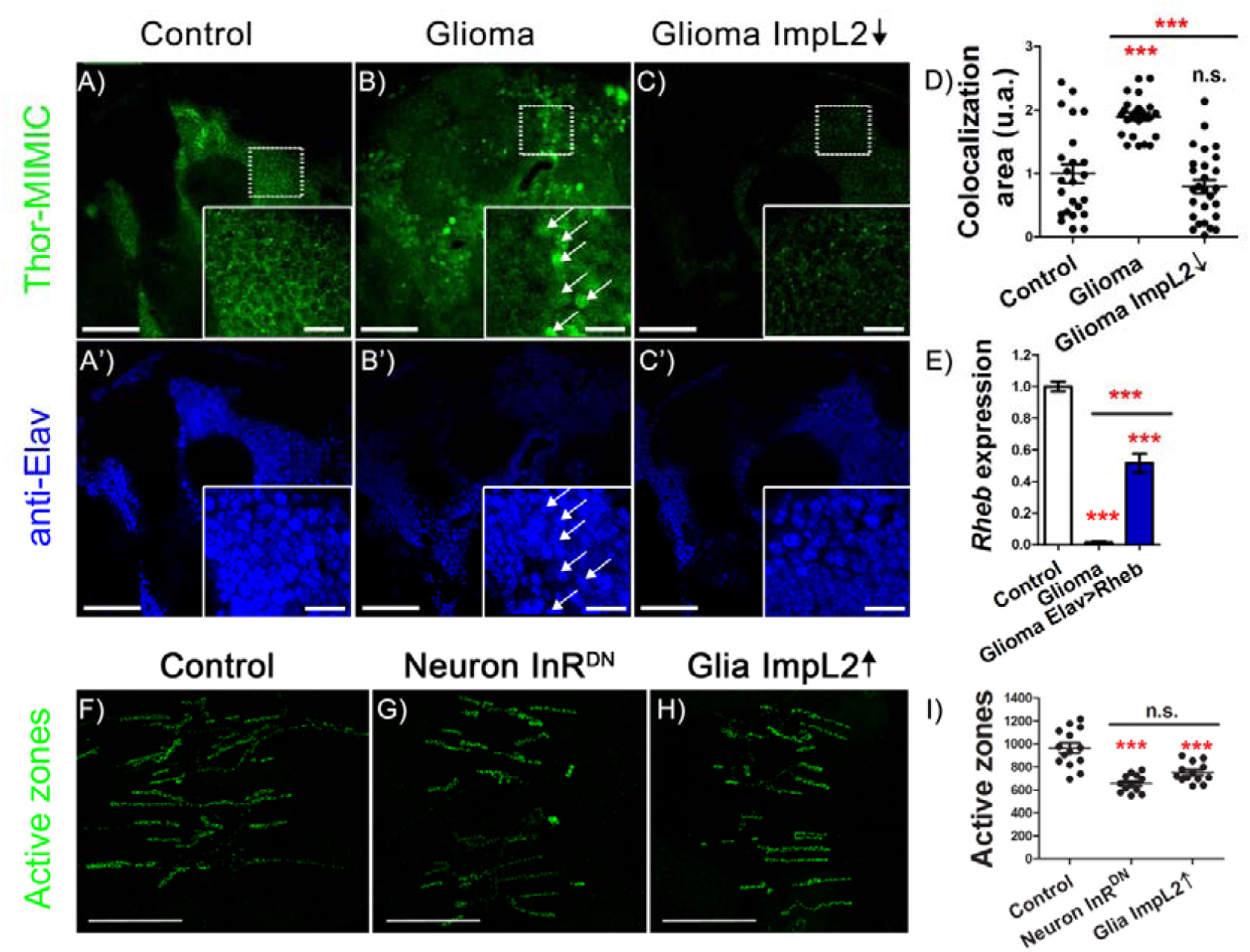
Glioma-secreted ImpL2 inhibits insulin pathway activity in neurons. Confocal microscopy images of adult brain of 7-day-old flies from A) *repo>UAS-LacZ*, B) *repo>UAS-dEGFR*^λ^, *UAS-dp110CAAX* and C) *repo>UAS-dEGFR*^λ^, *UAS-dp110CAAX, UAS-ImpL2* genotypes together with a Thor-MIMIC transgene (showed in green). Neurons are marked in blue (A’-C’). D) Quantification and statistical analysis of correlation area in *N* = 9 per genotype (ANOVA, post-hoc Bonferroni) (scale bar, 50 μm/10 μm). E) RT-qPCR of *Rheb* expression is downregulated in gliomas and the ectopical expression in neurons rescues the phenotype (ANOVA, post-hoc Bonferroni) in at least *N* = 30 per genotype. Adult neuromuscular junction (NMJ) from F) *D42>UAS-LacZ*, G) *D42>UAS-InR*^*DN*^ and H) *repo>UAS-ImpL2* animals after 7 days at 29ºC (active zones shown in green). *D42* is expressed in motor neurons. I) Quantification and statistical analysis of active zones in at least *N* = 20 per genotype (ANOVA, post-hoc Bonferroni) (scale bar, 50 μm) (\*\*\**p*-value>0,0001).

To further analyze insulin-dependent TOR activity we used a *THOR-MIMIC* line. The gene *thor* encodes for a protein that is involved in translational control. It is regulated by TOR and its expression can hence be used as a surrogate of TOR activity. In normal conditions, *thor* transcription remains at low but detectable levels (45). However, when Insulin activity is compromised, *thor* is highly transcribed, as reflected by *LacZ*- or *MIMIC*-*THOR* lines (46). Neurons confronted with GB cells have reduced Insulin signaling, as shown by *THOR-MIMIC* GFP expression. This effect on Insulin pathway is restored by down-regulating *ImpL2* in GB cells (fig 4A-D). All these results together suggest that *ImpL2* up-regulation in GB cells mediates the decreased insulin pathway activity detected in neurons.

However, the central function for insulin signaling pathway related to synaptogenesis was described mainly in larval NMJ synapsis (31). Whether or not insulin signaling plays a similar role in the central adult nervous system has not been evaluated yet. To do so, we expressed a dominant negative form of the *insulin receptor (InR)* in motor neurons and quantified the number of synapses. The number of synapses was reduced when compared to the control (fig 4F-I). Moreover, high levels of secreted ImpL2 from glial cells also shows a reduction in the synapse number, thus suggesting that the ImpL2 effect on synapsis is due to a deregulation of the insulin signaling in neurons.

### Insulin signaling in neurons mediates neurodegeneration and mortality in GB

Our results suggest that in a GB ImpL2 overexpression causes an effective decrease in neuronal Insulin pathway activity which in turn induces neurodegeneration. If this is true, restoring Insulin signaling specifically in neurons should prevent GB-induced neurodegeneration. However, in order to manipulate Insulin pathway activity in neuronal population simultaneously with GB induction in glial cells, we needed to use the *LexA/LexAOp* system (see Materials and Methods). We overexpressed *LexAOp*-d*Rheb* (thus activating Insulin signaling) in neurons using an *elav-LexA* line. d*Rheb mRNA* levels increase significantly upon LexA/LexAop system activation, therefore validating the LexAop-*Rheb* tool (fig 4E). The quantifications of adult NMJs show an increase in the synapse number in NMJ when compared to GB and even to control genotypes, thus showing a similar effect (although slightly stronger) to GB with low ImpL2 levels (fig 5A-C, G). In conclusion, the larval synaptogenic pathway regulated by Insulin signaling members is conserved in adult brains, at least for *InR* and *dRheb*.

**Fig 5.**
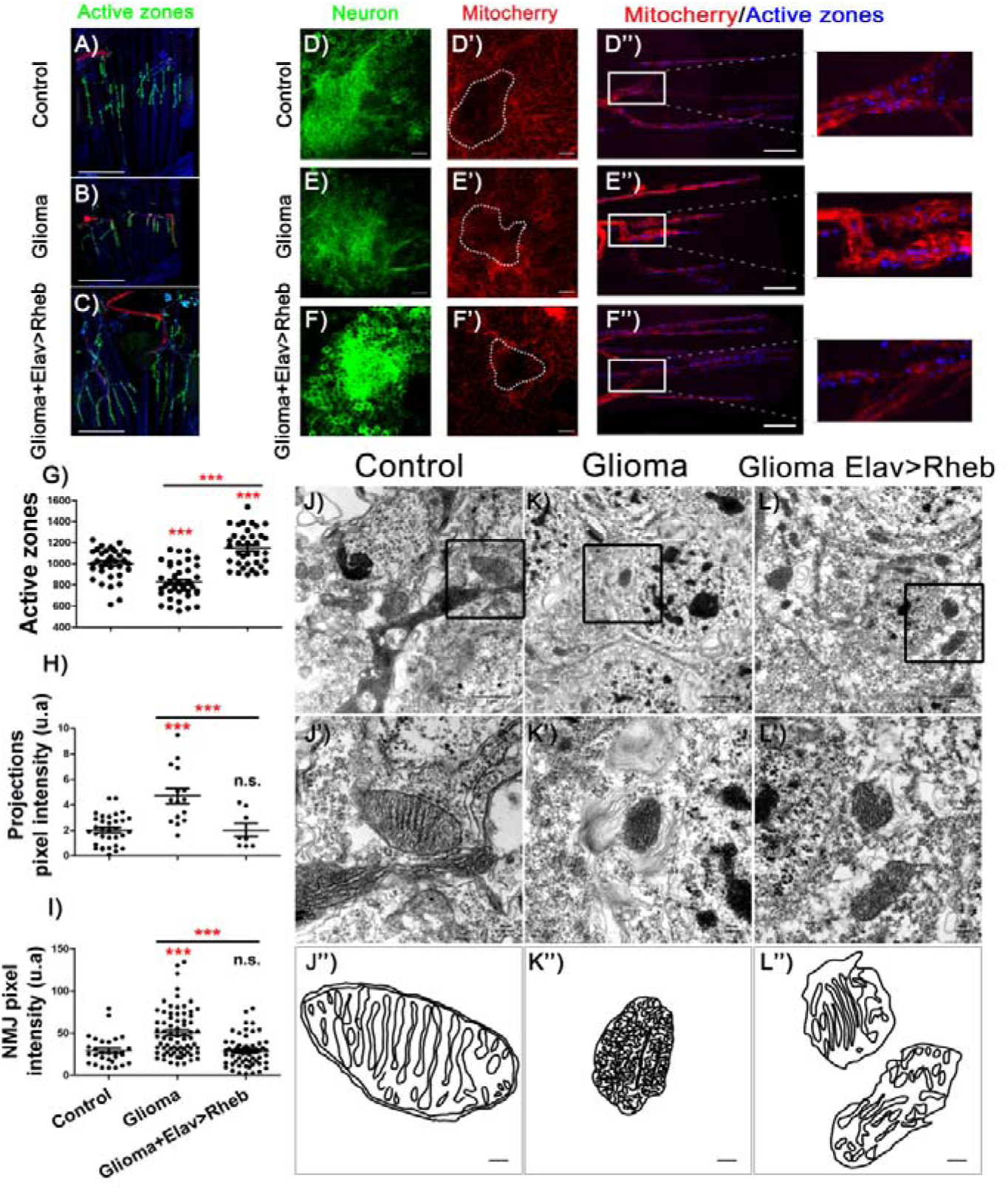
Glioma-induced mitochondrial alterations are rescued by *Rheb* overexpression. Adult neuromuscular junction (NMJ) from A) *repo>UAS-LacZ* (Control), B) *repo>UAS-dEGFR*^*λ*^, *UAS-dp110CAAX* (Glioma) and C) *repo>UAS-dEGFR*^*λ*^, *UAS-dp110CAAX*; *elav-LexA, LexAOp-Rheb* (Glioma+Elav>Rheb) after 7 days at 29ºC (active zones are marked in green). G) Quantification and statistical analysis of active zones number in at least *N* = 20 per genotype (ANOVA, post-hoc Bonferroni) (scale bar, 50 μm). Confocal microscopy images of adult brain and NMJ from D) *repo>UAS-LacZ*, E) *repo>UAS-dEGFR*^*λ*^, *UAS-dp110CAAX* and F) *repo>UAS-dEGFR*^*λ*^, *UAS-dp110CAAX*; *elav-LexA, LexAOp-Rheb* after 7 days at 29ºC (neural membrane shown in green), D’-F’) mitochondrial membrane in red and D”-F”) active zones in blue (scale bar, 10 μm). Quantification and statistical analysis of pixel intensity in H) projections and I) NMJ in at least *N* = 15 and *N* = 3 respectively per genotype (ANOVA, post-hoc Bonferroni). Electron microscopy images of neurons from brains of adult flies of J) *repo>UAS-LacZ*, K) *repo>UAS-dEGFR*^λ^, *UAS-dp110CAAX* and L) *repo>UAS-dEGFR*^λ^, *UAS-dp110CAAX; elav-LexA, LexAOp-Rheb* and a higher magnification of the area inside the black square showing the mitochondria in detail (J’-L’) (scale bar, 1 μm/ 100 nm respectively). J”-L”) Schematic representation of the ultrastructure of the cristae (scale bar, 100 nm).

Insulin signaling mediates glucose metabolism and mitochondrial physiology (47). Additionally, mitochondrial alterations are related to synapse dysfunction and neurodegeneration (48) (49). In particular, the transport of mitochondria through the axons is altered in other neurodegenerative diseases (50). Therefore, we investigated if GB progression affects neuronal mitochondria.

We used a *lexAop-mitocherry* reporter transgene to quantify the distribution of mitochondria in axon terminals. Pixel intensity quantification showed that GB causes a significant increase of mitochondria in the neuronal projections of Kenyon cells in the mushroom body (fig 5 D-F, D’-F’, H), compatible with a neurodegenerative process (51). In line with this, we observed an increase of mitochondria accumulated in NMJ boutons that correlates with lower number of synaptic boutons (5D”-F”, I). Both effects in neuronal projections and NMJs are prevented upon Insulin pathway signaling activation in the neuronal population of a GB brain (fig 5F,F’,F”). Consistently, electron microscopy images of neuronal mitochondria in GB samples show defects in the cristae, a typical feature of non-functional mitochondria (52, 53). This mitochondrial defective morphology is also reverted upon d*Rheb* overexpression in neurons (fig 5 J-L). Altogether, these results indicate that despite the higher number of mitochondria, neurons show dysfunctional mitochondria as a consequence of low insulin signaling. Increasing insulin pathway activity in neurons exposed to GB may be sufficient to recover functional organelles, as suggested by the synapse number.

GB brains with high insulin signaling in neurons also show a reduction in the number of glial cells and in tumor volume (fig 6 A-E). More importantly, GB causes premature death, an effect that is significantly rescued by overexpressing d*Rheb* in neurons (fig 6 F). These results suggest that restoration of neuronal insulin pathway activity improves the lifespan in animals with GB, thus linking synaptogenesis to a slower disease progression and providing a possible novel therapeutic target.

**Fig 6.**
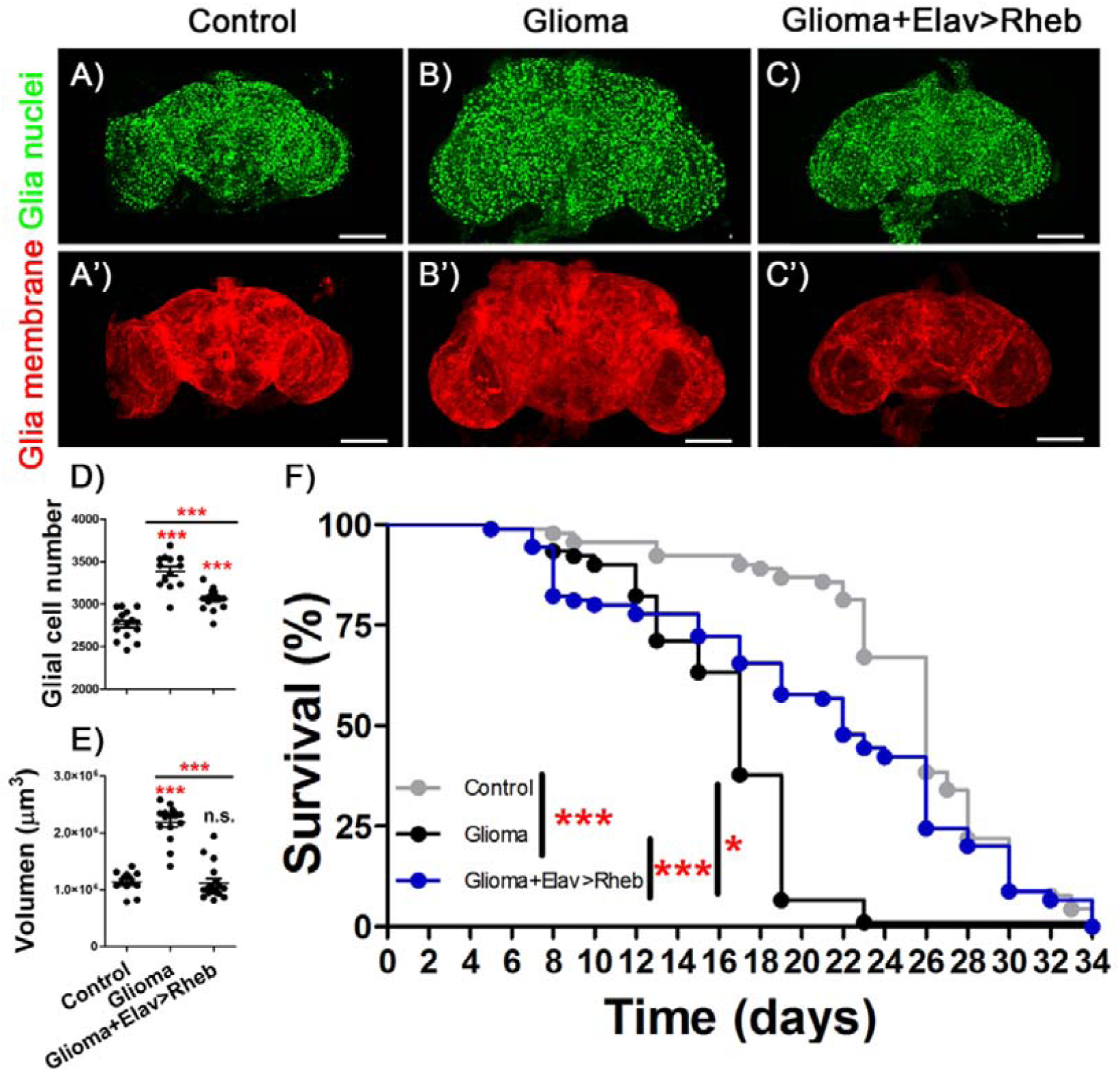
Overexpression of *Rheb* in neurons is protective against glioma effects. Confocal microscopy images of adult brains from A) *repo>UAS-LacZ*, B) *repo>UAS-dEGFR*^λ^, *UAS-dp110CAAX* and C) *repo>UAS-dEGFR*^λ^, *UAS-dp110CAAX; elav-LexA, LexAOp-Rheb* after 7 days at 29ºC with glial nuclei marked in green. A’-C’) Glial membrane is shown in red. D) Glial cell number quantification and E) glial membrane volume quantification for at least N = 13 per genotype (ANOVA, post-hoc Bonferroni) (scale bar, 100 μm). F) Graph shows a survival assay of *repo>UAS-LacZ* (grey), *repo>UAS-dEGFR*^λ^, *UAS-dp110CAAX* (black) and *repo>UAS-dEGFR*^λ^, *UAS-dp110CAAX, elav-LexA, LexAOp-Rheb* (blue) flies and statistical analysis in *N* = 90 (Mantel-Cox test) (\**p*-value>0,05,\*\**p*-value>0,005, \*\*\**p*-value>0,0001).

## DISCUSSION

GB is the most aggressive type of brain tumor. During GB progression tumoral cells extend a network of membrane projections that contribute to brain infiltration and results in poor prognosis for the patient. GB courses with a neurological decay that includes drowsiness, sleep disturbances, speech difficulties and other typical symptoms of neurodegeneration (3). For decades the origin of this decay was attributed to the high pressure caused by the GB solid mass and the associated edema. Despite the solid mass of the GB being removed after surgery, the neurodegenerative process continues, more likely due to the diffuse GB progression. This indicates that mechanisms underlying neurological decay are not restricted to the intracranial pressure and edema.

Recent publications suggested an active communication between GB cells and the surrounding healthy tissue, including neurons. Experiments done with human GB cells in mice xenografts revealed a physical interaction between GB cells and neurons as electrical and chemical synapses (54, 55). In this case neurons act as presynaptic structures whereas GB tumoral cells are postsynaptic elements. These so-called “synapses” are required for GB progression. Besides, we have recently described cellular mechanisms for GB to deplete Wingless (Wg)/WNT from neurons. GB cells project TMs that enwrap neurons and accumulate Frizzled1 (Fz1) receptor to vampirize Wg from neighboring neurons. This unidirectional mechanism facilitates GB proliferation and cause a loss of synapses in the neurons (17). Finally, here we also describe a one-way communication system from GB cells towards neurons. ImpL2 protein is originated in GB cells and target healthy neurons, but not vice-versa. ImpL2 binds InR ligands and act as an antagonist of the pathway. In consequence, insulin signaling is reduced in neurons which in turn cause mitochondrial aberrations, synapse loss and lethality (fig 7). Neuronal insulin signaling can be restored via *Rheb* upregulation, and this is sufficient to extend the lifespan of GB animals. These results suggest that reducing insulin signaling (and the subsequent neurodegeneration) is critical for GB proliferation, progression and invasion, and ultimately for the lethality caused by GB.

**Fig 7.**
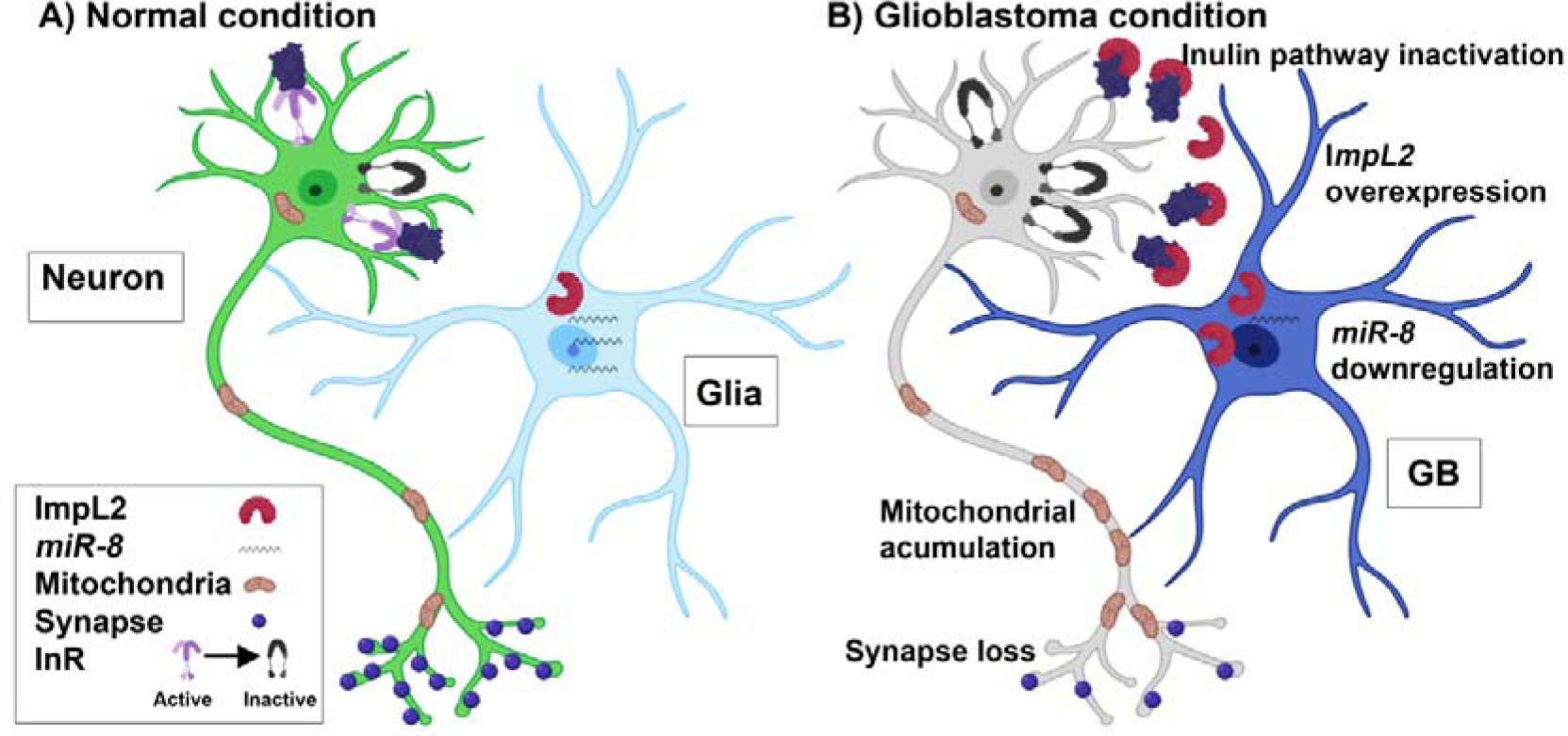
Schematic representation of GB effect over neuronal insulin pathway. A) In a physiological condition glial cells shows normal *miR-8* and ImpL2 expression levels. B) In a GB condition, *miR-8* downregulates its expression and ImpL2 levels increase, which in turn causes inactivation of the insulin pathway. As a consequence, low insulin signaling triggers different neuronal changes such as synapse loss and mitochondrial accumulation through the axon. The strategy used to revert the tumoral phenotype consist in the restoration of insulin signaling levels in the neuronal population either by overexpressing *Rheb* in neurons or upregulating/down-regulating *miR-8/ImpL2* expression in GB cells.

The regulation of *ImpL2* expression in GB cells seems to depend on microRNAs, at least in part. In cancer, micro-RNAs have emerged as general regulators key for tumoral progression, including GB (20). Interestingly, *miR-200* family (*miR-8* in *Drosophila*) play a key role in human GB. We have described the presence and relevance of *miR-8* in GB progression as a regulator of *ImpL2* expression. However, there are no predicted *miR-8* binding sites in *ImpL2* sequence; therefore, it is unlikely that *miR-8* regulates directly *ImpL2 mRNA* stability. One possibility is the existence of mediator proteins that depend directly on *miR-8*. The most evident possibility is a transcription factor whose mRNA stability is sensitive to *miR-8* and acts as a transcriptional regulator of *ImpL2.* However, the regulation of *ImpL2* and the association to microRNAs is a matter for future studies. Another intriguing point is the differential effects that both *miR-8* overexpression and *ImpL2* down-regulation have on GB growth. Whereas an excess of *miR-8*, that in turn reduces *ImpL2*, is unable to reduce GB cell number, direct downregulation of *Impl-2* significantly reduces the growth of GB cells. Nevertheless, it is known that most of microRNAs control several mRNAs, thus *miR-8* overexpression might affect to different extent other *mRNAs* than just *ImpL2* mRNA, which may account for such differences.

Altogether, our data suggest that the progression of brain tumors in *Drosophila* depends not only in the intrinsic properties of the tumoral cells, but also in the physiological condition of the surrounding cells (fig 7). GB patients respond differently to the progression of the GB: some patients survive for a few months whereas others survive for years. If we accept that the coordinated effect in GB and neurons result in differential tumor progression, the vast differences in how patients respond to GB could be, in part, dependent on genetic or epigenetic conditions related to InR signaling genes in neurons, and probably other pathways such as WNT or Hedgehog.

## Materials and methods

### Fly stocks and genetics

All fly stocks were maintained at 25°C (unless otherwise specified) on a 12/12h light/dark cycles at constant humidity in standard medium. The stocks used from Bloomington Stock Center were: *tub-Gal80*^*ts*^ (BL-7019), *Repo-Gal4* (BL-7415*), D42-Gal4* (BL-8816), *UAS-InR DN* (Bl-2852), *UAS*-*myr*-*RFP* (BL-7119) *UAS-LacZ* (BL-8529), *UAS-ImpL2RNAi* (BL-55855), *UAS-ImpL2MIMIC* (BL-59246), *lexAop-mitocherry* (BL-66530). Others fly stocks used were: *UAS-miR-8-sensor* (41), *UAS-miR-8-RFP* (56), *Elav-LexA* (BL52676), *lexAop-Rheb, UAS-HRP::CD2 (*gifted by L.Luo, Watts el al., 2004), *UAS-dEGFR*^λ^;*UAS-dp110*^*CAAX*^ (gift from R. Read, Read et al., 2009) *UAS-ImpL2* (gift from H. Stocker), LexAop-Rheb, (gift from Nuria Romero).

The glioma-inducing line contains the *UAS-dEGFR*^λ^, *UAS-dp110*^*CAAX*^ transgenes that encodes for the constitutively active forms of the human orthologues PI3K and EGFR respectively (16). *Repo-Gal4* line drives the *Gal4* expression in every glial cell (57) and combined with the *UAS-dEGFR*^λ^, *UAS-dp110*^*CAAX*^ line allow us to generate a glioma thanks to the Gal4 system (58). *Elav-LexA* line drives the expression to neurons (59), allowing us to manipulate neurons in a glioma combining *LexA* and *Gal4* expression systems (60).

Gal80^TS^ is a repressor of the Gal4 activity at 18°C, though at 29°C is inactivated (McGuire et al., 2003). The *tub-Gal80*^*ts*^ construct was used in all the crosses to avoid the lethality caused by the glioma development during the larval stage. The crosses were kept at 17°C until the adult flies emerged. To inactivate the Gal80^ts^ protein and activate the Gal4/UAS system to allow the expression of our genes of interest, the adult flies were maintained at 29°C for a period indicated in each experiment.

### Immunostaining and Image acquisition

Adult brains were dissected and fixed with 4% formaldehyde in phosphate-buffered saline for 20 minutes whereas adult NMJ were fixed 10 minutes; in both cases, samples were washed 3×15 min with PBS+0.4% triton, blocked for 1 h with BSA 1%, incubated overnight with primary antibodies, washed 3×15 min, incubated with secondary antibodies for 4 h and mounted in Vectashield mounting medium with DAPI. The primary antibodies used were anti-repo mouse (1/200, DSHB), anti-bruchpilot -NC82- mouse (1/50, DSHB), anti-HRP rabbit (1/400, Cell Signalling) anti-GFP rabbit (1:500, DSHB). The secondary antibodies used were Alexa 488 or 647 (1/500, Life Technologies). Images were taken by a Leica SP5 confocal microscopy. To quantify synapses and number of glial cells, we analyzed confocal images with the Imaris software (www.bitplane.com). It allows to select intensity points of a specific diameter that correspond either to the synapses (0.5 µm) or glial cell nuclei (2 µm). Using this software, it is also possible to determine the volume of the glial membrane. For the analysis of pixel intensity, the “measurement log” tool of Photoshop CS5 was used. For the analysis of co-localization rates, “co-localization” tool from LAS AF Lite software (Leica) was used.

### RNA extraction, reverse transcription and qPCR

For RNA extraction, 1- to 4-day-old male adults were entrained to a 12:12 h LD cycle for 7 days at 29°C and then collected on dry ice at ZT 6. Total RNA was extracted from 30 heads of adult males of the Control (*repo>LacZ*), Glioma (*repo> UAS-dEGFR*^λ^, *UAS-dp110*^*CAAX*^) and *repo> UAS-dEGFR*^λ^, *UAS-dp110*^*CAAX*^, *elav-LexA, LexAOp-Rheb* genotypes after 7 days of glioma development. RNA was extracted with TRIzol and phenol chloroform. Total RNA concentration was measured by using NanoDrop ND-1000. cDNA was synthetized from 1 mg of total RNA using M-MLV RT (Invitrogen). cDNA samples from 1:5 dilutions were used for real-time PCR reactions. Transcription levels were determined in a 14 mL volume in duplicate using SYBR Green (Applied Biosystem, Foster City, CA) and 7500qPCR (Thermo Fisher, Waltham, MA). We analysed transcription levels of *ImpL2, Rheb* and *Rp49* as housekeeping gene reference.

Sequences of primers were: RP49 F: GCATACAGGCCCAAGATCGT, Rp49 R: AACCGATGTTGGGCATCAGA, ImpL2 F: CCGAGATCACCTGGTTGAAT, ImpL2 R: AGGTATCGGCGGTATCCTTT, Rheb F: CGACGTAATGGGCAAGAAAT and Rheb R: CAAGACAACCGCTCTTCTCC.

After completing each real-time PCR run, outlier data were analysed using 7500 software (Applied Biosystems). Ct values of triplicates from 3 biological samples were analysed calculating 2DDCt and comparing the results using a t-test with GraphPad (GraphPad Software, La Jolla, CA).

### Viability and survival assays

Lifespan was determined under 12:12 h LD cycles at 29°C conditions. Three replicates of 30 1- to 4-day-old male adults were collected in vials containing standard *Drosophila* media and transferred every 2-3 days to fresh *Drosophila* media.

### Electron microscopy

Adult brains of *repo>LacZ, repo> UAS-dEGFR*^λ^, *UAS-dp110*^*CAAX*^ and *repo> UAS-dEGFR*^λ^, *UAS-dp110*^*CAAX*^, *elav-LexA, LexAOp-Rheb* animals expressing CD2-HRP in glial membranes were dissected after 7 days of glioma development and fixed with 4% formaldehyde in phosphate-buffered saline for 30 minutes. The samples were washed twice with PBS and incubate with R.T.U. VECTASTAIN kit (VECTOR) for 30 min at RT and washed once with PBS. Followed by an incubation in darkens with SIGMA FAST 3,3’-Diaminobenzidine Tablet SETS (SIGMA) for 75 min at RT, washed once with PBT and incubate with 4% formaldehyde + 2% glutaraldehyde for 1 hour and store at 4°C. Following fixation samples were washed three times in 0.1 m phosphate buffer. Postfixation was done in 1% osmium tetroxide + 1% potassium ferrocyanide for 1 hour at 4°C, three washes in H2O2dd and incubated in PBS 0.1M + 0.15% tanic acid for 1 min, washed once in PBS 0.1M and twice in H2O2dd. Following by incubation in 2% uranil acetate 1 hour at RT in darkness and three washes in H2O2dd. Dehydration was done in ethanol series (30%, 50%, 70%, 90% and 3×100%). The samples were infiltrated with increasing concentrations of epoxy resin TAAB-812 (TAAB Laboratories) in propilen oxid, encapsulated in BEEM capsules to polymerize 48 hours at 60°C. Ultrathin sections of 70-80 nm were cut using Ultracut E microtome (Leica) and stained with 2% uranyl acetate solution in water and lead Reynols citrate. The grids were examined with JeolJEM1400Flash (Tokyo, Japan) electron microscope at 80 kV. Images were taken with a OneView (4Kx4K) CMOS camera (Gatan).

### Statistics

The results were analyzed using the GraphPad Prism 5 software (www.graphpad.com). Quantitative parameters were divided into parametric and non parametic using the D’Agostino and Pearson omnibus normality test and the variances were analyzed with F test. Student’ t test and ANOVA test with Bonferroni’s post-hoc were used in parametric parameters, using Welch’s correction when necessary. To the non-parametric parameters, Mann-Whitney test and Kruskal-Wallis test with Dunns post-hoc were used. The survival assays were analyzed with Mantel-Cox test. The *p* limit value for rejecting the null hypothesis and considering the differences between cases as statistically significant was *p* <0.05 (*). Others *p* values are indicated as ** when p< 0.01 and *** when p<0.001.

